# The long and the short of it: unlocking nanopore long-read RNA sequencing data with short-read tools

**DOI:** 10.1101/2020.06.28.176727

**Authors:** Xueyi Dong, Luyi Tian, Quentin Gouil, Hasaru Kariyawasam, Shian Su, Ricardo De Paoli-Iseppi, Yair David Joseph Prawer, Michael B. Clark, Kelsey Breslin, Megan Iminitoff, Marnie E. Blewitt, Charity W. Law, Matthew E. Ritchie

## Abstract

Application of Oxford Nanopore Technologies’ long-read sequencing platform to transcriptomic analysis is increasing in popularity. However, such analysis can be challenging due to small library sizes and high sequence error, which decreases quantification accuracy and reduces power for statistical testing. Here, we report the analysis of two nanopore sequencing RNA-seq datasets with the goal of obtaining gene-level and isoform-level differential expression information. A dataset of synthetic, spliced, spike-in RNAs (“sequins”) as well as a mouse neural stem cell dataset from samples with a null mutation of the epigenetic regulator *Smchd1* were analysed using a mix of long-read specific tools for preprocessing together with established short-read RNA-seq methods. We used *limma-voom* to perform differential gene expression analysis, and the novel *FLAMES* pipeline to perform isoform identification and quantification, followed by *DRIMSeq* and *limma-diffSplice* (with *stageR*) to perform differential transcript usage analysis. We compared results from the sequins dataset to the ground truth, and results of the mouse dataset to a previous short-read study on equivalent samples. Overall, our work shows that transcriptomic analysis of long-read nanopore data using short-read software and methods that are already in wide use can yield meaningful results.

## Introduction

Short-read sequencing technology has underpinned transcriptomic profiling research over the past decade. The sequencing platforms offered by companies such as Illumina Inc. provide high read accuracy (>99.9%) and throughput which allows many samples to be profiled in parallel. One major limitation of short-read sequencing technology is the modest read lengths offered (currently up to 600 bases), which makes accurate isoform quantification and novel isoform discovery challenging. Long-read sequencing offers a distinct advantage in this regard, with the ability to generate reads that are typically in the 1-100 kilobase (kb) range(1), which spans the typical length distribution of spliced genes in human (for protein coding genes 1-3 kb is typical with outliers such as Titin at more than 80kb) thereby allowing the sequencing of entire isoforms. This however comes at the expense of lower throughput and reduced accuracy compared to short-read sequencing. The two main technology platforms that dominate the field of long-read sequencing are Pacific Biosciences’ Single-Molecule Real Time (SMRT) sequencing and Oxford Nanopore Technologies’ (ONT) nanopore sequencing.

Previous work on long-read transcriptomic data focuses on transcript-level analysis, especially in the discovery of novel isoforms(2–4). Some long-read specific methods have been developed for this task. Reference-based methods, such as *TALON*(5), compares reads to existing gene and transcript models to create novel models. Reference-free methods, such as *FLAIR*(6), maps reads to the reference genome, clusters alignments into groups and collapses them into isoforms. Differential transcript usage (DTU) is another transcript-level analysis that is of great interest(6–8). DTU analyses examine differences in the relative proportions of expressed isoforms between two conditions. The *DRIMSeq*(9) method performs DTU analysis on transcript-level RNA-seq counts using a Dirichlet-multinomial model. Alternatively, tools developed for differential exon usage analysis, such as the *diffSplice* function(10) in *limma* and *edgeR* packages have also been adapted to DTU analyses(11). Both *DRIMSeq* and *diffSplice* methods were developed for short read data. The *stageR* package(12) can be used to control the false discovery rate (FDR) of DTU analyses through its stage-wise method which screens potential DTU genes using gene-level *p*-values before selecting the transcripts with evidence of DTU.

Previous studies have looked at gene-level analysis of nanopore data but study design limited the methods used. Soneson *et al.*(13) concluded that read coverage in native RNA libraries (~0.5 million aligned reads per flow cell) were too low for gene-level analyses, resulting in low power and high variability. Li *et al.*(7) worked around this by simply using fold-changes to identify differentially expressed genes for three ONT MinION direct RNA *Caenorhabditis elegans* samples, however the lack of statistical testing could lead to unreliable results. Jenjaroenpun *et al.*(14) used *DESeq2*(15) to perform differential expression analysis on direct RNA transcript-level counts, but gene-level expression was not studied.

In this study, we performed gene- and isoform-level analyses of two nanopore long-read transcriptome sequencing datasets that follow a simple replicated experimental design: a synthetic “sequins”(16) PCR-cDNA dataset, and a mouse neural stem cell direct-cDNA dataset. We obtained meaningful results using an analysis pipeline that mostly comprised of *“off-the-shelf”* methods developed for short-read data, despite our datasets having only a few million long reads per sample. We found our results for the common two-group experimental design to be reliable in that they are broadly consistent with the available ground-truth or findings from a previous short-read experiment. We found existing methods for isoform identification from long-read data to be unreliable, and introduce a novel method, *FLAMES*, as part of our isoform-level analysis pipeline.

## Materials and methods

### Study design

Mouse neural stem cells (NSCs) from 4 wild type (WT) and 3 MommeD1 mutated (*Smchd1*-null)(17) samples were prepared and sequenced, together with 3 “other” samples from a different experiment. Samples were sequenced in two batches, each containing 6 samples. One WT and one “other” sample were sequenced in both batches as technical replicates.

Technical replicates of synthetic “sequin” RNA standards(16) from two mixes (A and B) were prepared and sequenced. These samples contain the same transcripts but at variable molar ratios to simulate biological differences in gene expression and alternative splicing. Among the 76 synthetic genes, 21 were up-regulated and 23 were down-regulated in mix B compared to mix A. The corresponding transcripts of 28 genes were expressed at different proportions between the two mixes, resulting in DTU for 62 out of 160 transcripts.

### Biological materials

Synthetic “sequin” RNA standards were obtained from the Garvan Institute of Medical Research.

NSCs were derived as described in Chen *et al.*(18). Cells were grown in NeuroCult Stem Cell medium (StemCell Technologies #05702) with cytokines: NeuroCult NSC Basal Medium (Mouse) (StemCell Technologies #05700) supplemented with NeuroCult Proliferation Supplement (Mouse) (StemCell Technologies #05701), 0.25 mg/mL rh EGF (StemCell Technologies #02633) and 0.25 mg/mL rh bFGF (StemCell Technologies #02634). We extracted total RNA with Trizol and purified polyA RNA with the NEBNext Poly(A) mRNA Magnetic Isolation Module (E7490).

### Nanopore sequencing and data preprocessing

Sequin cDNA libraries were constructed with SQK-PCS109 cDNA-PCR sequencing and SQK-PBK004 PCR Barcoding kits using the supplied protocol. Briefly, duplicate libraries of each mix (A1, A2, B1, and B2) were constructed using 15 ng as input for cDNA synthesis. Samples were barcoded 1 to 4 using the supplied PCR barcodes. Transcripts were amplified by 14 cycles of PCR with a 6-minute extension time.

Sequencing libraries were individually purified using Beckman Coulter 0.8x AMPure XP beads and quantified using an Invitrogen Qubit 4.0 Fluorometer (ThermoFisher Scientific). Equimolar amounts of each sample were pooled to a total of approximately 160 fmol (assuming median transcript size is 1 kb), and quality control of the pooled library was performed using Agilent Technologies TapeStation 4200. The final library was loaded onto an R9.4.1 MinION flow cell and sequenced for 65 hours with a buffer refuel at 24 hours (using 250 mL buffer FB) using the ONT GridION platform. The *fast5* files were base-called by *Guppy* version 3.2.8 using configuration file *dna_r9.4.1_450bps_hac.cfg* to obtain `fastq` files. *MinKNOW* version 3.6.0 was used to trim adaptor sequences and demultiplex barcoded reads. Both *Guppy* and *MinKNOW* are only available to ONT customers via the community site (https://community.nanoporetech.com/).

For the NSC dataset we prepared direct-cDNA libraries from 40 ng purified polyadenylated RNA. We combined the ONT direct-cDNA sequencing (SQK-DCS108) protocol (version DCB_9036_v108_revG_30Jun2017) with the one-pot native barcoding protocol(19) with extended incubation times (using SQK-LSK109 and EXP-NBD103 kits) for library preparation of the first batch, and used the updated kits SQK-DCS109 and EXP-NBD114 for the second batch (protocol PDCB_9093_v109_revA_04Feb2019). We loaded 100 ng of the final libraries on one PromethION flow cell (FLO-PRO002) per batch. The `fast5` files were base-called by *Guppy* version 3.1.5 using configuration file *dna_r9.4.1_450bps_hac_prom.cfg* to yield `fastq` files. We used *Porechop*(20) to trim adaptor sequences from reads and demultiplex barcoded reads. The “other” samples were removed in downstream analysis. For an overview of our analysis pipeline see Figure 1A.

**Fig. 1.**
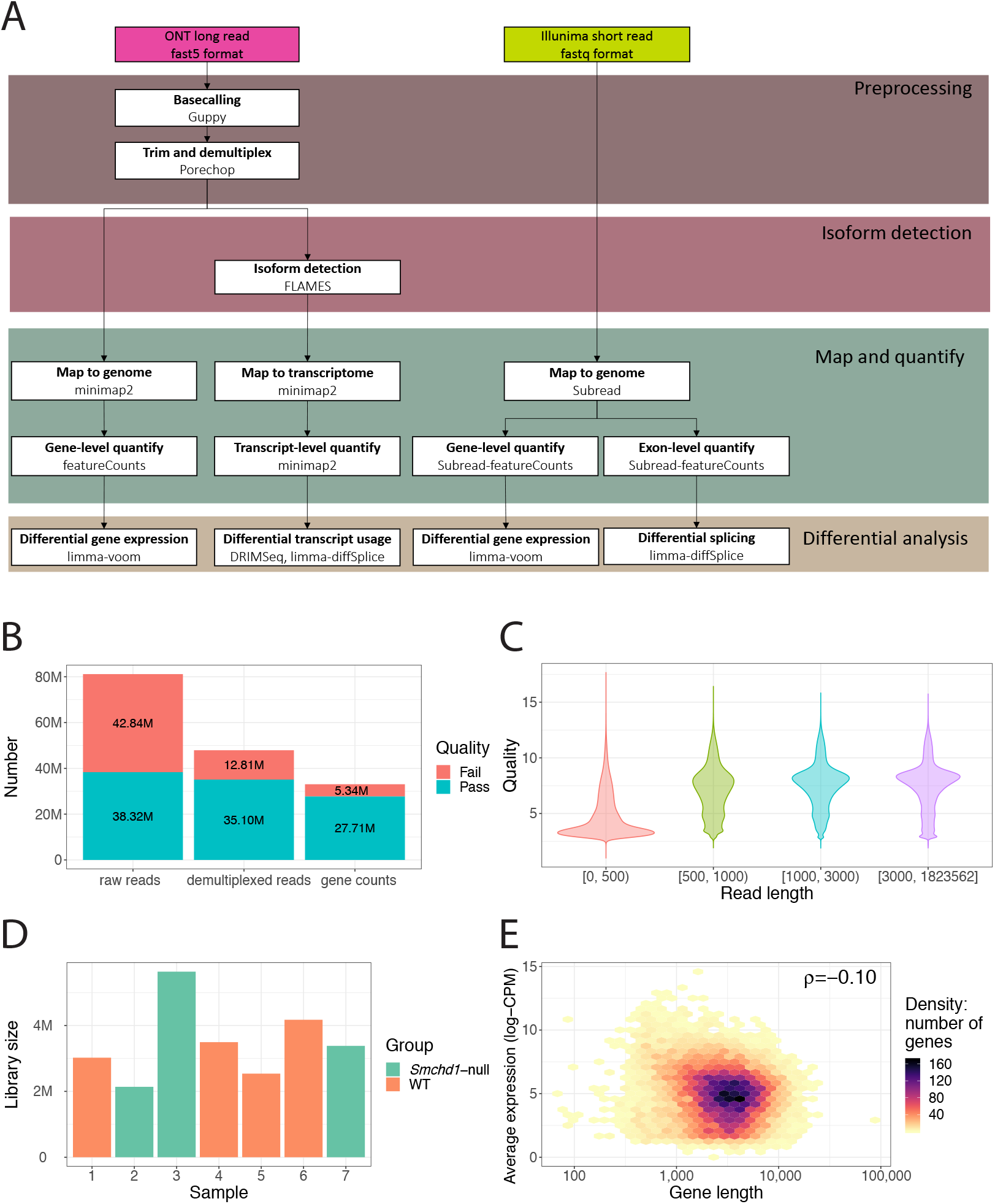
Analysis workflow and quality metrics. **(A)** Overview of the analysis workflow used to process the mouse NSC direct-cDNA long-read and short-read RNA-seq data. **(B)** The number of raw reads, trimmed and demultiplexed reads and gene-level counts in the NSC dataset, stratified by average base quality score (Pass: >7, turquoise; Fail: ≤7, red). **(C)** Distribution of read quality in the NSC dataset, stratified by read length. Read quality is defined by the average base quality score of a read. **(D)** The total number of reads assigned to each sample in the NSC dataset (green: *Smchd1*-null samples; orange: WT samples). **(E)** A hexagonal 2D density plot showing the correlation between gene length and average gene expression (log-CPM) in the NSC dataset.

### Genomic alignment

The NSC reads were aligned to mouse *mm10* genome using *minimap2* version 2.17-r943-dirty(21) to get `bam` files. The genome alignments were performed with the arguments *-ax splice -uf -k14 −junc-bed*, allowing spliced alignments on the forward transcript strand to map with higher sensitivity. It also uses annotated splice junctions to improve the accuracy of mapping at junctions. Gencode release M23 (GRCm38.p6) annotation(22) was used to provide information on known splicing junctions. The sequins ONT reads were mapped to the artificial chromosome *chr*IS_R using *minimap2* with the arguments *-ax splice −MD*. The bam files were sorted and indexed using *samtools* version 1.6(23).

### Gene abundance estimation

Mapped reads were assigned to individual genes and counted by the *feature-Counts*(24) function in the R(25)/Bioconductor(26) package *Rsubread* version 1.34.4(27, 28). We used the in-built *mm10* annotation for the NSC data, and sequins annotation GTF file version 2.4 for the sequins data. Arguments *isLongRead=TRUE* and *primaryOnly=TRUE* were used to indicate that the input data contains long reads and to count primary alignments only.

### Differential gene expression analysis

Genes in the NSC dataset were annotated using R/Bioconductor package *Mus.musculus*(29), and read counts from technical replicates were combined. For both datasets, we organized and preprocessed the count data using the R/Bioconductor package *edgeR* version 3.26.8(30, 31). Lowly expressed genes were removed using the *filterByExpr* function with default arguments. Normalization factors were calculated using the trimmed mean of M-values method(32). Differential gene expression (DGE) analysis was performed using the *limma-voom* pipeline version 3.40.6(10, 33, 34), with sample-specific quality weights(35). Linear models were fitted with either genotype or sequin mix information to create the design matrix, followed by empirical Bayes moderation of *t*-statistics(36). Raw *p*-values were adjusted for multiple testing(37).

### Mouse NSC short-read data

We obtained DGE results from a previous Illumina short-read RNA-seq study comparing mouse *Smchd1*-null and WT NSC samples(18, 38) available at http://bioinf.wehi.edu.au/folders/smchd1/ and from GEO (accession number GSE65747). Using a *limma-voom* pipeline, the study reported 1,197 differentially expressed (DE) genes (adjusted *p*-value cutoff of 0.01). We further restricted this list (adjusted *p*-value cutoff of 0.0001) to give us 218 up- and 54 down-regulated genes in *Smchd1*-null samples when compared to WT samples. This cutoff resulted in similar numbers of significant genes between the short- and long-read datasets. The DE genes were compared to that of the NSC long-read data using ROAST gene set testing(39) with 9,999 rotations.

### Transcript-level analysis

We used three different tools to perform isoform detection and quantification: *FLAIR* version 1.5.0(6) and *TALON* version 5.0.0(5) for the sequins data, and *FLAMES* version 0.1.0 for both sequins and NSC datasets. Default parameters were used to run *FLAIR*. *TranscriptClean*(40) version 1.02 which performs reference-based error correction was applied prior to running *TALON* version 5.0. Transcripts identified by *TALON* were filtered using default setting.

*FLAMES* (short for ‘**F**ull-**L**ength tr**A**nscript quantification, **M**utation and **S**plicing analysis’) is a novel method and software tool developed for long-read RNA-seq data, available at https://github.com/LuyiTian/FLAMES. It requires sorted bam files with reads aligned to the genome as input. Scanning each gene, *FLAMES* groups reads that are similar as isoforms (transcript start/end site differs by <100bp and splice site differs by <10bp). The isoforms are then compared against a reference annotation (Gencode release M23 (GRCm38.p6)(22)), and transcript sequences are extracted from the isoform assembly. All reads are then re-aligned to both the known and assembled transcripts and quantified. High-confidence isoforms are those identified with at least 10 supporting reads.

After running each isoform detection tool, *SQANTI*(41) was used to classify identified isoforms by comparing them to the annotation. Lowly expressed transcripts were removed from downstream analysis. We kept transcripts with 10 or more counts in at least 3 samples in the NSC data, and in at least 2 samples in the sequins data. For both datasets, genes in every sample were also required to have an associated gene count (obtained by summing counts across all transcripts for a given gene) of 10 or more.

DTU analysis was performed using two methods: the R/Bioconductor package *DRIMSeq* version 1.12.0(9), and *diffSplice* from the *limma* package version 3.40.6. Originally, *DRIMSeq* was designed for use with transcript-level counts in short-read data, giving adjusted *p*-values at both the gene-level and feature-level (transcripts). *DiffSplice* analyses exon-level counts in short-read data to indirectly call for differences in isoform proportions, and reports adjusted *p*-values at the gene-level (Simes adjustment and/or F-tests) and exon-level (*t*-tests). For long-read data, we applied the *diffSplice* to transcript-level counts rather than exon-level counts as carried out by Love *et al.*(11). Additionally, the stage-wise method from R/Bioconductor package *stageR* version 1.6.0(12) was also applied to the raw *p*-values from *DRIMSeq* (gene- and transcript-level) and *diffSplice* (Simes and *t*-tests) methods for FDR control to give *stageR* gene- and transcript-level adjusted *p*-values.

### Data and code availability

RNA-seq data can be accessed from Gene Expression Omnibus (GEO) under accession numbers GSE151984 (sequins) and GSE151841 (NSC data). All code used to perform these analyses are available from https://github.com/XueyiDong/LongReadRNA.

## Results

### Data quality

To assess the quality of our long-read datasets, raw long reads were pre-processed and assigned to gene-level counts using an appropriate reference genome. Figure 1B shows the number of reads (or amount of information) retained after some crucial steps in processing the NSC data. A total of ~81 million reads were successfully sequenced and base-called. A relatively small proportion of those, ~60% or ~48 million reads, were detected with adaptor and barcode sequences in the trimming and demultiplexing steps. Most of the reads that did not pass these steps were of low quality and marked as “fail” reads (average base quality score ≤7). The reads were then mapped to the genome and assigned to genes, producing ~28 million gene-level counts. For the sequins dataset, ~7.3 million raw “pass” reads yielded ~5.7 million gene-level counts (Supplementary Figure S1).

In the NSC dataset, median read length is about 800 bases, where low quality “fail” reads are generally shorter (median of ~300 bases). Figure 1C shows that short reads (< 500 bases) tend to have low read quality relative to longer reads, similar to that observed in Soneson *et al.*(13). The quality of “extra long” reads (≥3000 bases) were similar to that of “long” (1000 to 2999 bases) and “medium” (500 to 999 bases) length categories, indicating Nanopore’s ability to detect transcripts in this size range. A small proportion (< 1%) of reads exceed 5 kilobases. Similar to the NSC dataset, the sequins dataset had a median read length of 996 bases, which is sightly longer than its expected value of 908 bases.

The library size (sum of gene counts) of samples in the NSC dataset varied between 2.1 million to 5.6 million reads (Figure 1D). As comparison, the library size of samples in the short-read NSC study(18) were ranging between 18.6 million to 23.2 million reads.

### Gene expression analysis

In short-read RNA-seq, transcripts (or genes) are fragmented for sequencing, such that longer transcripts can be over-represented relative to transcripts that are shorter. As a consequence, DGE analyses are biased towards the detection of genes (or transcripts) that are relatively long(42). Also, DGE analyses may be confounded by DTU, such that gene-level counts are affected by the varying proportions of transcripts with varying lengths. One advantage to long-read RNA-seq protocols is that they do not include the fragmentation step, and should theoretically be unbiased to gene length. To examine this, we looked at the relationship between gene length and gene expression using log_2_-counts-per-million (log-CPM) values. Gene length is weakly associated with expression in both long-read datasets; Pearson correlation of 0.10 for the NSC dataset (Figure 1E), and −0.05 for sequins dataset (Supplementary Figure S2). Whereas, correlation in the short-read NSC study(18) is greater, at 0.20.

We applied the *limma-voom* workflow designed for short-read DGE analysis to our long-read data. We first analysed the sequins data to check whether the approach was appropriate using the ground-truth available. The analysis was carried out using *voomWithQualityWeights* to account for sample-level heterogeneity by estimating sample-specific quality weights based on similarity of gene expression within the same group. The sample weights were combined with *voom* precision weights that are based on the mean-variance relationship estimated from the data. Even though there were only 69 genes present in the dataset (as opposed to tens of thousands in a typical dataset), the mean-variance trend observed for the sequins data (Supplementary Figure S3) was similar to that of short-read RNA-seq data(33).

Linear modelling on the gene-level counts were carried to obtain estimated log_2_fold-change (logFC) values between mix A and B. Estimated values were highly correlated (R_2_=0.933) with expected logFCs (Figure 2A). Using an adjusted *p*-value cutoff of 0.05, 21 down-regulated and 18 up-regulated genes were detected between mix B and A. There were no false discoveries, and only 2 of the truly differentially expressed genes were not detected. Overall, results from the sequin synthetic control data indicate that the *limma-voom* pipeline is powerful and reliable when applied to long-read data, so we applied it to the NSC dataset also.

**Fig. 2.**
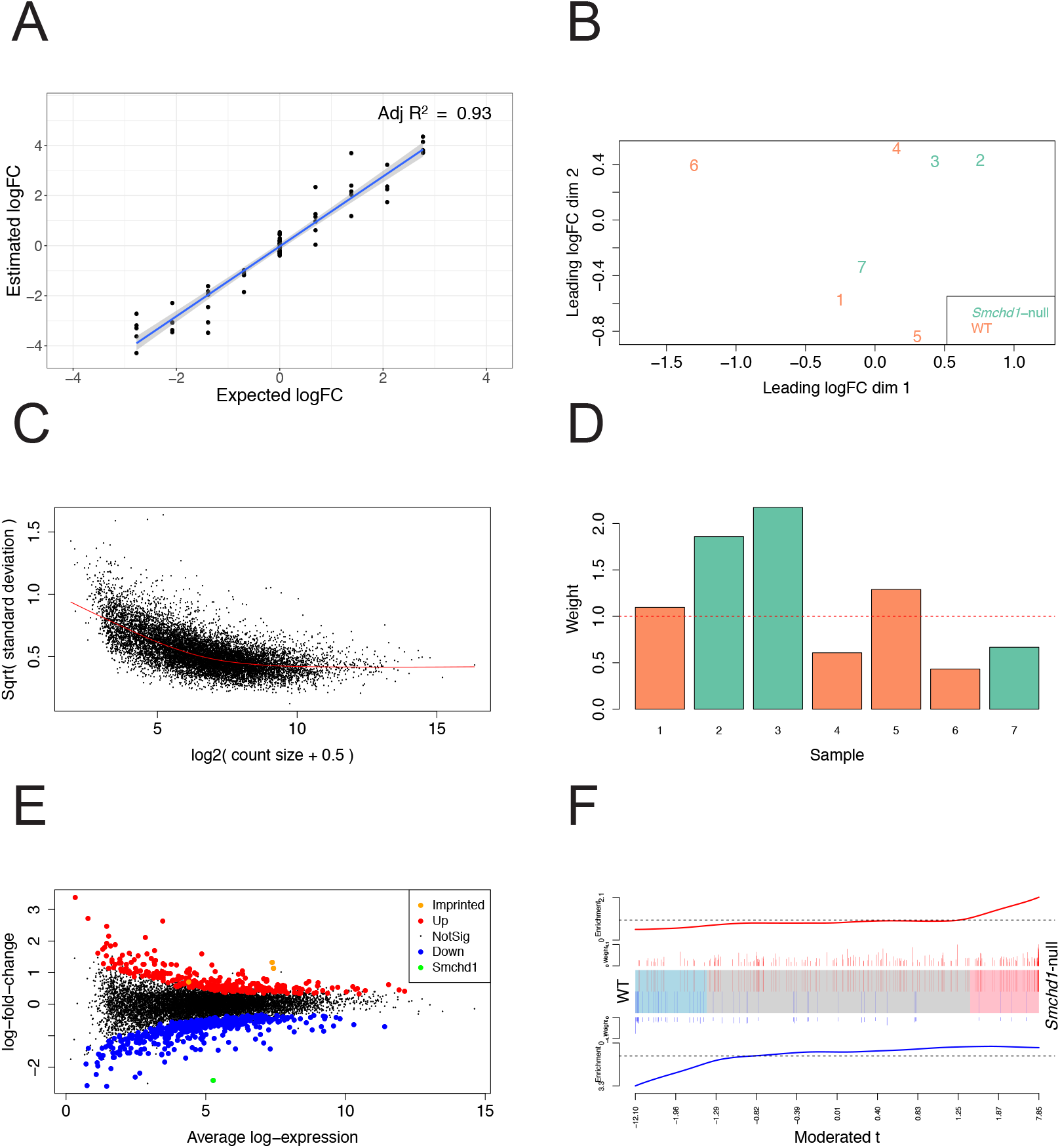
Results for differential gene expression analysis. **(A)** Correlation between the observed log2 FC between mix A and B and the expected log2 FC in the sequins data. The blue line is the linear regression line. **(B)** MDS plot showing the relationship between NSC samples based on gene-level logCPM. **(C)** Voom mean-variance trend in NSC data where points represent genes, and **(D)** sample-specific weights obtained from the *limma-voom* workflow (colour-coded by genotype). **(E)** Gene-level plot of logFC for *Smchd1*-null versus WT plotted against average log2 - expression values. Differentially expressed genes are highlighted (red: up-regulated genes, blue: down-regulated genes). **(F)** The barcode plot shows the correlation between our long-read differential expression results and the results from a previous short-read dataset collected on the same NSC sample types. Each vertical bar represents a DE gene from the previous short-read study (red: up-regulated genes, blue: down-regulated genes), and the position of the bar on the x-axis represents the moderated t-statistic of the same gene in our long-read results. The length of the vertical lines represent the logFC of the gene in the short-read results. The red worm on the top and the blue worm at the bottom represent the relative enrichment of the vertical bars in each part of the plot.

Unsupervised clustering by multidimensional scaling (MDS) was used to observe the relationships between NSC samples. Dimension 1 in the MDS plot roughly separates samples by genotype (Figure 2B), although a *Smchd1*-null sample (sample 7) is positioned more closely to WT samples.

The mean-variance trend for this dataset is strikingly similar to that which is typically observed in short-read RNA-seq experiments(33) (Figure 2C); even more so than in the sequins dataset. Estimated sample weights favoured samples that distinguished groups across dimension 2 of the MDS plot, giving samples 2 and 3 in the *Smchd1*-null group weights that are greater than 1, as well as samples 1 and 5 in the WT group (Figure 2D). Using the default adjusted *p*-value cutoff of 0.05, only 17 genes were detected as DE. Using a more liberal adjusted *p*-value cutoff of 0.25 to account for the small library sizes, detected 413 down-regulated and 321 up-regulated genes between *Smchd1*-null and WT samples (Figure 2E). The *Smchd1* gene, which was depleted in *Smchd1*-null samples, was detected as the most significantly down-regulated gene in the comparison (highlighted in Figure 2E) and serves as a positive control for our analysis. In a previous short-read study on the same mouse NSC groups(18, 38), the imprinted genes *Ndn*, *Mkrn3* and *Peg12* were reported as up-regulated. These genes were also found to be DE in the long-read dataset (highlighted in Figure 2E). Further comparison between the short- and long-read datasets was carried out using a barcode plot (Figure 2F). The barcode plot shows that most of the genes that were up-regulated in the short-read dataset (red vertical lines in the plot) also tend to be up-regulated in our long-read dataset (positioned towards the right of the plot). Specifically, the genes that were most highly up-regulated in the short-read dataset as ranked by logFC (long red vertical lines), are also highly up-regulated in the long-read data (further right in the plot). The same goes for down-regulated genes in the short-read dataset (blue vertical lines in the plot), which tend to be down-regulated in the long-read dataset (positioned towards the left of the plot). We tested concordance of the datasets formally by applying the ROAST gene set testing method(39) to our long-read data. Using both up- and down-regulated gene sets from the short-read dataset, weighted on their logFC values, ROAST returned an “up” *p*-value of 0.10, which indicates that transcriptional changes for the comparison of *Smchd1*-null versus WT are somewhat consistent between the two datasets (up-regulated genes in the short-read data tend to be up-regulated in the long-read data, and down-regulated genes in the long-read data tend to be down-regulated in the long-read data). The relatively large ROAST *p*-value and overall lack of power to detect differentially expressed genes is likely due to relatively low sequencing levels per sample and within-genotype sample heterogeneity in the long-read experiment.

### Transcript-level analysis

Transcript-level analysis of nanopore RNA-seq data usually starts with isoform detection. To test which tool is best suited to nanopore data, we compared two popular tools, *FLAIR* and *TALON* with our novel *FLAMES* pipeline on the sequins dataset. Ideally, all transcripts that appear in the sequins annotation should be detected, and there should not be any novel isoforms. Our results showed that *FLAMES* detected the most sequin transcripts (Figure 3A, ‘full splice match’ category) and the fewest artefactual isoforms (Figure 3A, other categories). While most sequin transcripts were also detected by *FLAIR*, a disproportionately large number of artefactual isoforms were also identified, especially those classified as ‘novel in catalog’ for which we know there are none. *TALON* only detected about half of the sequin transcripts, and many antisense isoforms. When we looked into the number of reads assigned to transcripts in each category (Figure 3B), the majority of counts in *FLAMES* were from known isoforms, while about half of the counts in *FLAIR* were from artefactual isoforms. The total number of read counts from *FLAMES* (~4M) and *FLAIR* (~2.4M) are similar, while for *TALON* it was much lower (~4.4K). Results from the sequins dataset indicated that the novel *FLAMES* pipeline outperformed the other two methods.

**Fig. 3.**
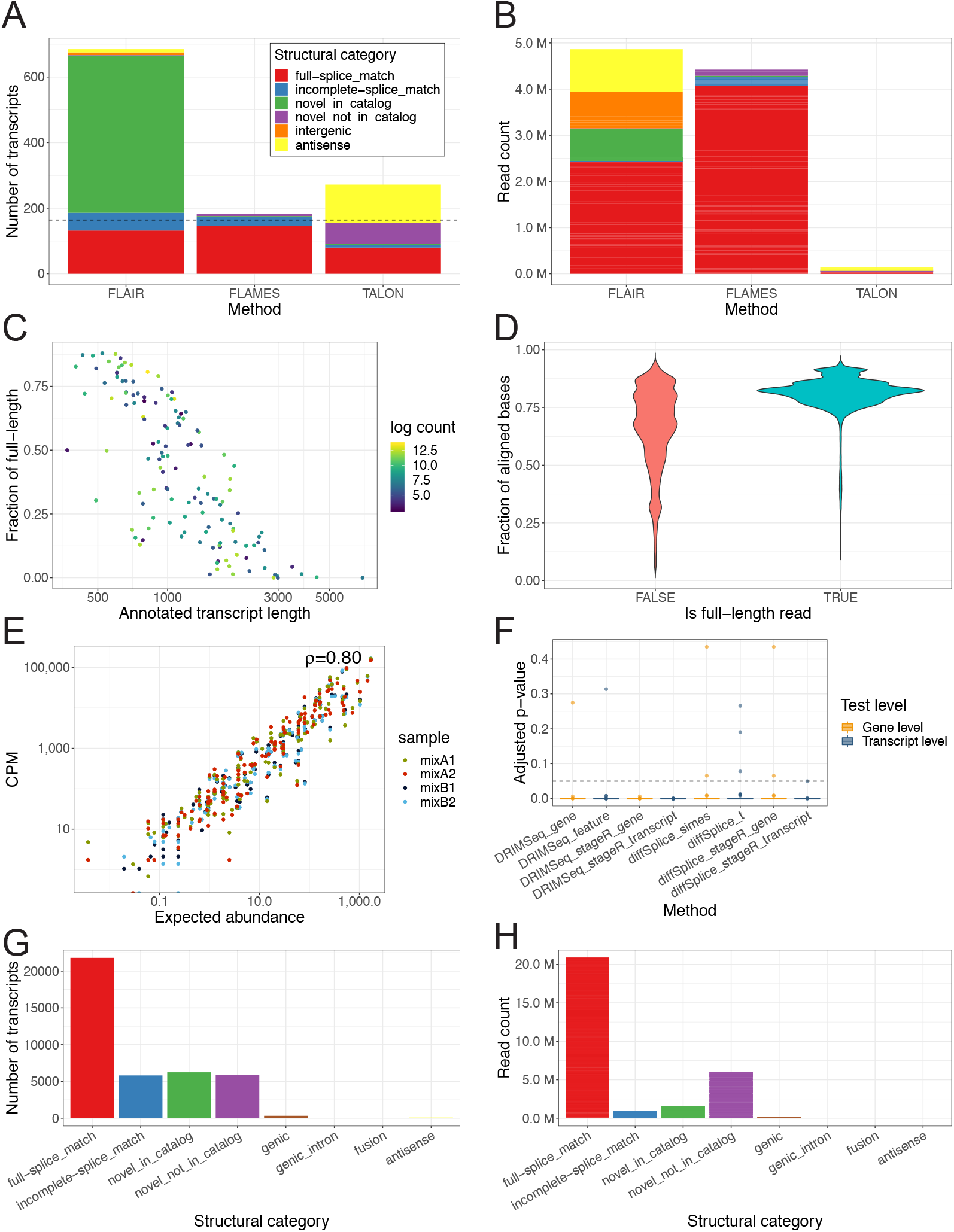
Isoform identification and differential transcript usage analysis. **(A)** A bar plot showing the number of discovered isoform types in the sequins dataset. The bars are separated into isoform categories (by colour), and the dashed line represents the true number of isoform types. **(B)** A bar plot showing the number of counts from isoforms in the sequins dataset. The bars are separated into isoform categories (by colour) from which the counts are associated with. **(C)** A scatter plot showing the correlation between the fraction of full-length reads assigned to a transcript and the length of the annotated transcript. Dots are coloured by log2 count of transcripts. **(D)** A violin plot showing the aligned fraction of reads (calculated as the aligned length divided by read length), stratified by whether the read is full-length. **(E)** The correlation between observed transcript counts and expected transcript abundance of each gene from each sequins sample. **(F)** A box plot showing the distribution of adjusted *p*-values from different tests of DTU for the true DTU genes or transcripts in sequins data. The dashed horizontal line shows the adjusted *p*-value cutoff of 0.05. Boxes are coloured by whether the test is performed on gene level or transcript level. **(G)** A barplot showing the number of discovered isoforms in each category output by *FLAMES* in the NSC dataset. **(H)** A barplot showing the number of counts from isoforms in each category output by *FLAMES* in the NSC dataset.

To further assess the performance of *FLAMES* and the quality of the dataset, we calculated the coverage fraction of transcripts by individual reads. Here, reads covering 95% or more of the bases of their corresponding transcript are defined as “full-length”. In our sequins data, 48% of reads were found to be full-length. Reads assigned to longer transcripts are less likely to be full-length (Figure 3C), consistent with findings from Jenjaroenpun *et al.*(14). We also observed that full-length reads have a higher fraction of aligned bases (Figure 3D). Looking further into the isoforms identified by *FLAMES*, we found that ‘full splice match’ isoforms tend to be longer than those in the other categories (Supplementary Figure S4), but they tend to miss some bases at the transcription start sites and/or transcription termination sites. This suggests that some reads may be truncated in our sequins dataset, which presumably occurs during library preparation of sequencing.

Next, we adapted *DRIMSeq* and the *diffSplice* function in *limma* for DTU analysis of long-read data. We also combined the methods with the stage-wise analysis from the *stageR*(12) package since it was recommended in the *DRIM-Seq* vignette for statistical improvement and enhanced biological interpretation of results. We expect good performance of DTU analyses comparing mixes A and B since the observed CPM values of sequin transcripts were highly correlated with their expected abundances (Figure 3E). Indeed, all of the DTU methods performed well, with no false discoveries for any of the methods (using an adjusted *p*-value cutoff of 0.05) (Figure 3F, Supplementary Figure S5). The true positive rate (TPR) (detecting genes/transcripts as having DTU when they truly have DTU) was very high for all of the methods (TPR>0.89), with *DRIMSeq* slightly out-performing *diffSplice* (Supplementary Figure S6). *StageR* transcript-level testing further improved the results of both *DRIMSeq* and *diffSplice* (TPR=1). *StageR* gene-level testing also improved the results from *DRIMSeq*, but not for *diffSplice*. We demonstrate that our pipeline and combination of methods for transcript-level analysis produces accurate transcript quantification and identification of DTU, and provides confidence for application to other long-read transcriptomic datasets.

We then applied our transcript-level analysis workflow to the NSC dataset. The *FLAMES* pipeline returned 40,221 unique isoforms from 10,986 genes, of which 48% were classified as novel (Figure 3G), which is a lot more than what was observed in the sequins dataset. Since we observed a very low level of falsely discovered isoforms in the sequins data, we assume that that most of these novel isoforms are real. This suggests that the current mouse transcript annotation is incomplete. Of the mapped reads, 29.4% were assigned to novel isoforms, the majority of which were from the ‘novel not in catalog’ category (Figure 3H). Both DTU methods (*DRIMSeq* and *diffSplice*) found only one gene *Pisd* as having DTU for *Smchd1*-null versus WT using an adjusted *p*-value cutoff of 0.25. *Pisd* was not DE at the gene-level (adjusted *p*-value=0.44). Transcript *ENSMUST00000201980.4* in *Pisd* was identified to have differential usage between the two groups (Supplementary Figure S7 and S8).

Using *limma-diffSplice* applied to exon counts, we also performed differential exon usage analysis on the NSC short-read data and found 87 significant genes (adjusted *p*-value cutoff of 0.05). The results, however, were discordant between short- and long-read datasets, with only 3 common genes in the top 200 most significant genes identified by *limma-diffSplice* for both datasets. *Pisd* was not identified to have significant differential splicing in the short-read analysis (adjusted *p*-value=1.00).

## Discussion

Our DGE analysis uses a *limma-voom* workflow and shows that results from PCR-cDNA and direct-cDNA long-reads are reliable, such that estimated results are comparable to the known truth in the sequins synthetic control dataset, and concordant with that of a previous short-read study for the NSCs. Although the total library size in the sequins dataset is lower than that of the NSC dataset, more reads were assigned per gene since the dataset contains a small set of genes, which improved power for DGE analysis. Overall, comparisons using long-read experiments suffer from a lack of statistical power due to low library sizes. It would be desirable for long-read transcriptomic studies to have total read numbers that are more comparable to what is routinely achieved in short-read experiments (20-50M reads per sample is not unusual). We expect this to occur in the near future as throughput of long-read experiments increases.

We also looked into transcript-level analysis of long-read data and found our novel *FLAMES* pipeline to be reliable in both isoform detection and quantification. The high false positive rate of *FLAIR* for isoform detection suggests that its reference-free algorithm needs further improvement to adapt to high error rates in long-read sequencing. Despite methods being designed originally for short-read data, *DRIMSeq* and *diffSplice* (in combination with *stageR* or not) performed very well in DTU analyses of our sequins data using transcript-level counts. We believe these methods could be applied to other datasets with confidence, but may lack power to detect DTU genes if transcript counts are very low. A potential issue in the application of the methods to our NSC dataset is that altered expression of the gene (*Smchd1*) may not affect RNA splicing mechanisms. Discordance between our results for short- and long-read NSC studies may be a reflection of the differences in exon-level counting in short reads versus transcript-level counting in long reads, rather than inconsistencies in the *diffSplice* method itself. Notably, relative to DGE analyses, a DTU analysis further splits gene-level counts into associated isoforms which reduces power for statistical testing. For this reason, the power to detect DTU genes in the NSC long-read dataset is reduced relative to the sequins dataset since the latter contains far fewer expressed genes and transcripts to begin with, such that transcripts have higher counts on average.

Our study is the first to test a pipeline for gene-level DGE analysis and transcript-level DTU analysis of nanopore long-read RNA-seq data. Whilst the sequencing depth is relatively low, we are still able to obtain reasonable results using pre-existing methods designed for short reads, namely the *limma* and *DRIMSeq* software. We expect that other short-read tools such as *edgeR* and *DESeq2* may also be appropriate, although this has not been tested. Exploring the strengths and weaknesses of different analysis methods on data arising from both the Nanopore and PacBio long-read platforms using a specially designed benchmarking dataset is planned as future work.

We hope that our analysis will encourage further research into the potential for long-read RNA-seq to be used in place of short-read RNA-seq, allowing for the simultaneous exploration of gene-level and isoform-level changes within the same experiment in a more comprehensive way.

## ACKNOWLEDGEMENTS

This project was supported by National Health and Medical Research Council (NHMRC) Project Grant (GNT1098290 to MEB and MER), a Career Development Fellowship (GNT1104924 to MER) an Early Career Fellowship (GNT1072662 to MBC), a Bellberry-Viertel Senior Medical Research Fellowship (to MEB), a Melbourne Research Scholarship to XD and LT and Victorian State Government Operational Infrastructure Support and Australian Government NHMRC IRIISS.

---

**Fig. S1.**
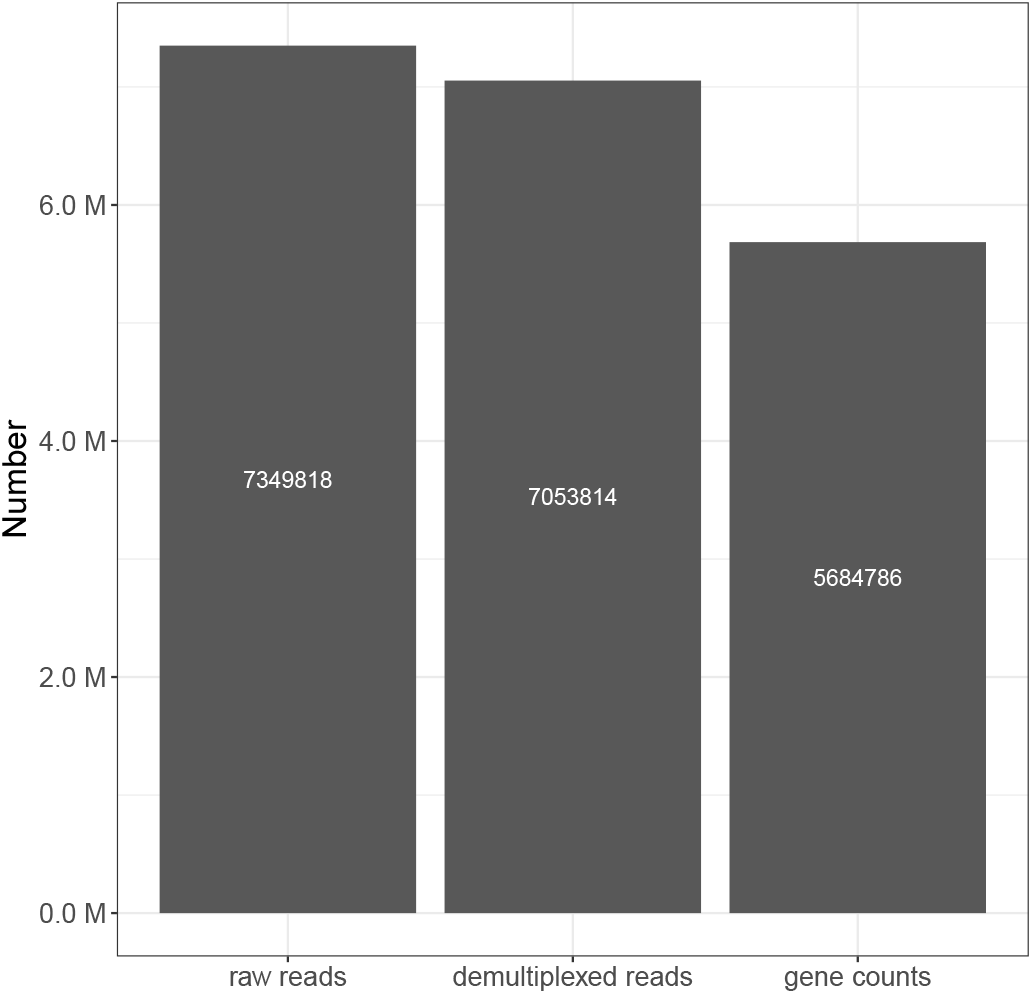
The number of pass (average base quality score >7) raw reads, trimmed and demultiplexed reads and assigned reads in the sequins dataset.

**Fig. S2.**
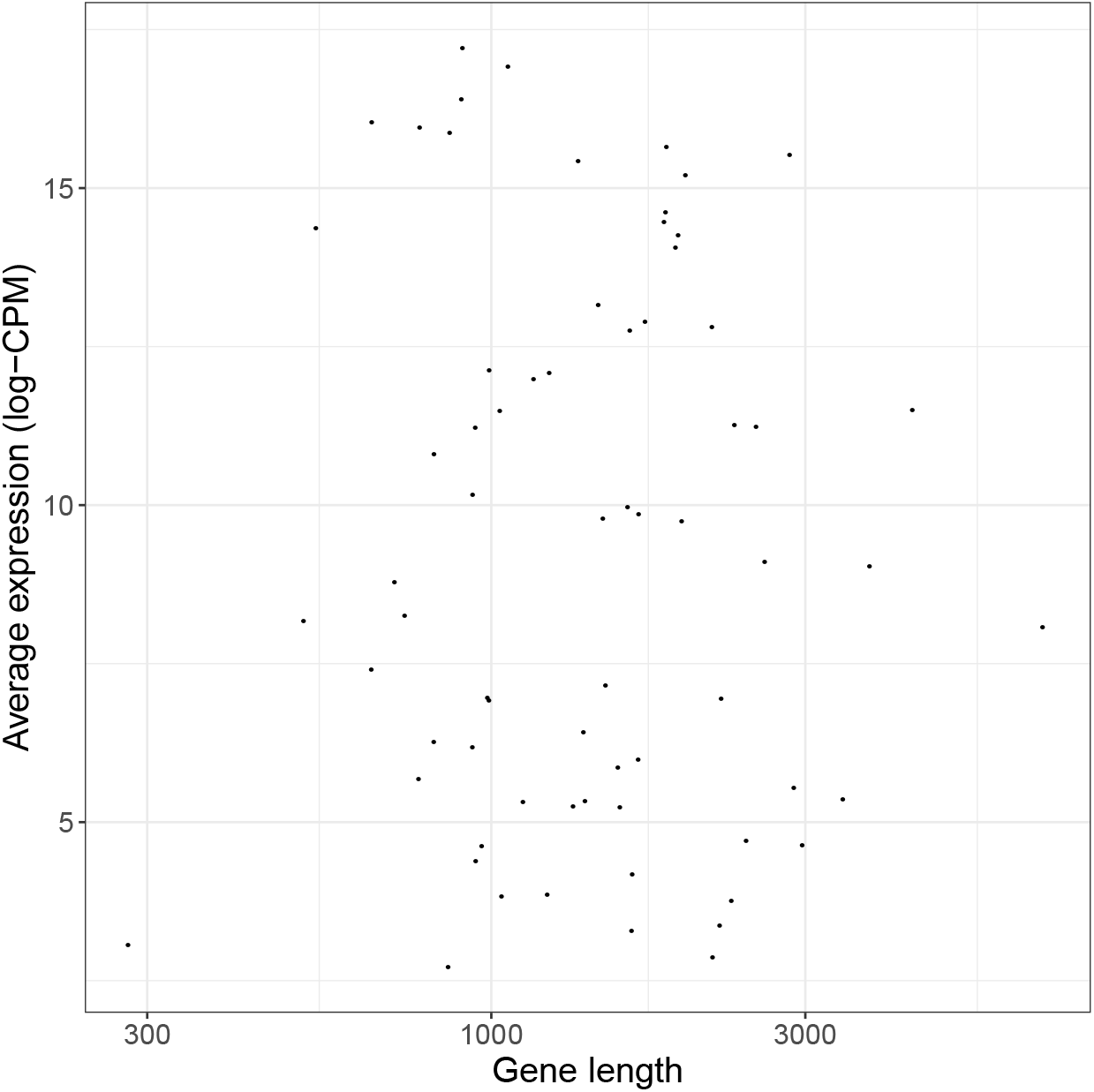
Correlation between gene length and average gene expression (log-CPM) in the sequins dataset.

**Fig. S3.**
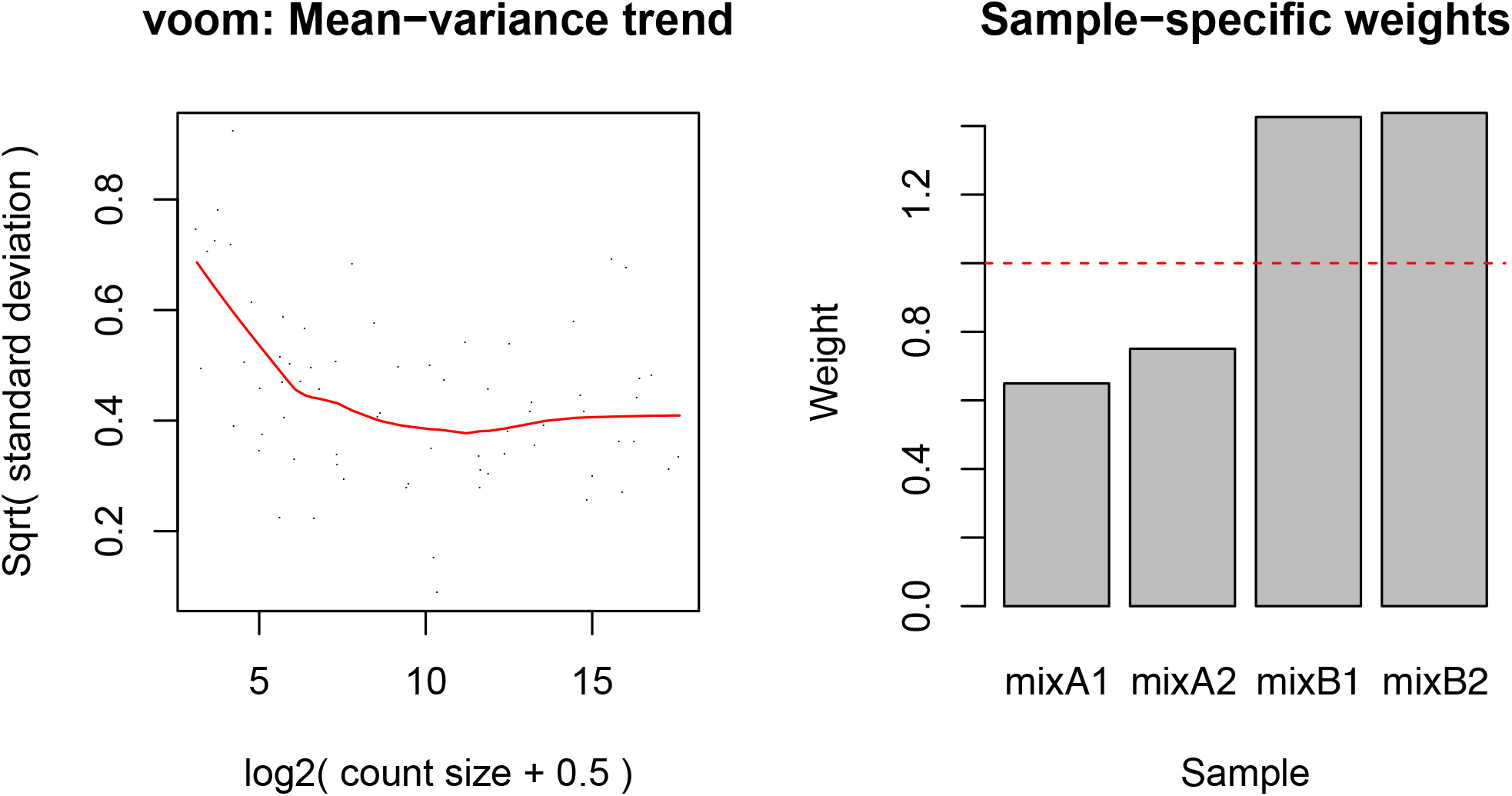
Voom mean-variance trend in the sequins data where points represent genes, and sample-specific weights obtained from the *voomWithQualityWeights* function.

**Fig. S4.**
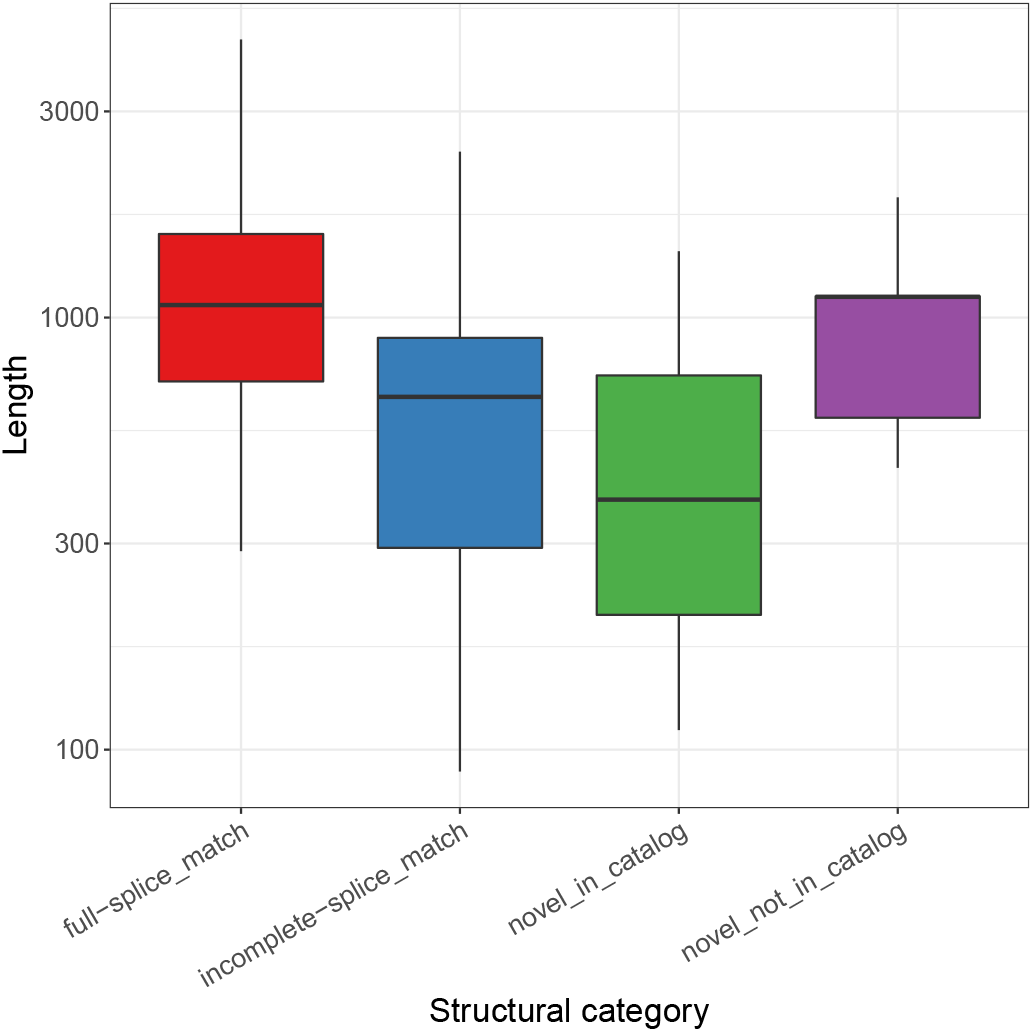
A box plot showing the length distribution of isoforms identified by *FLAMES* in the sequins dataset, stratified by isoform structural categories.

**Fig. S5.**
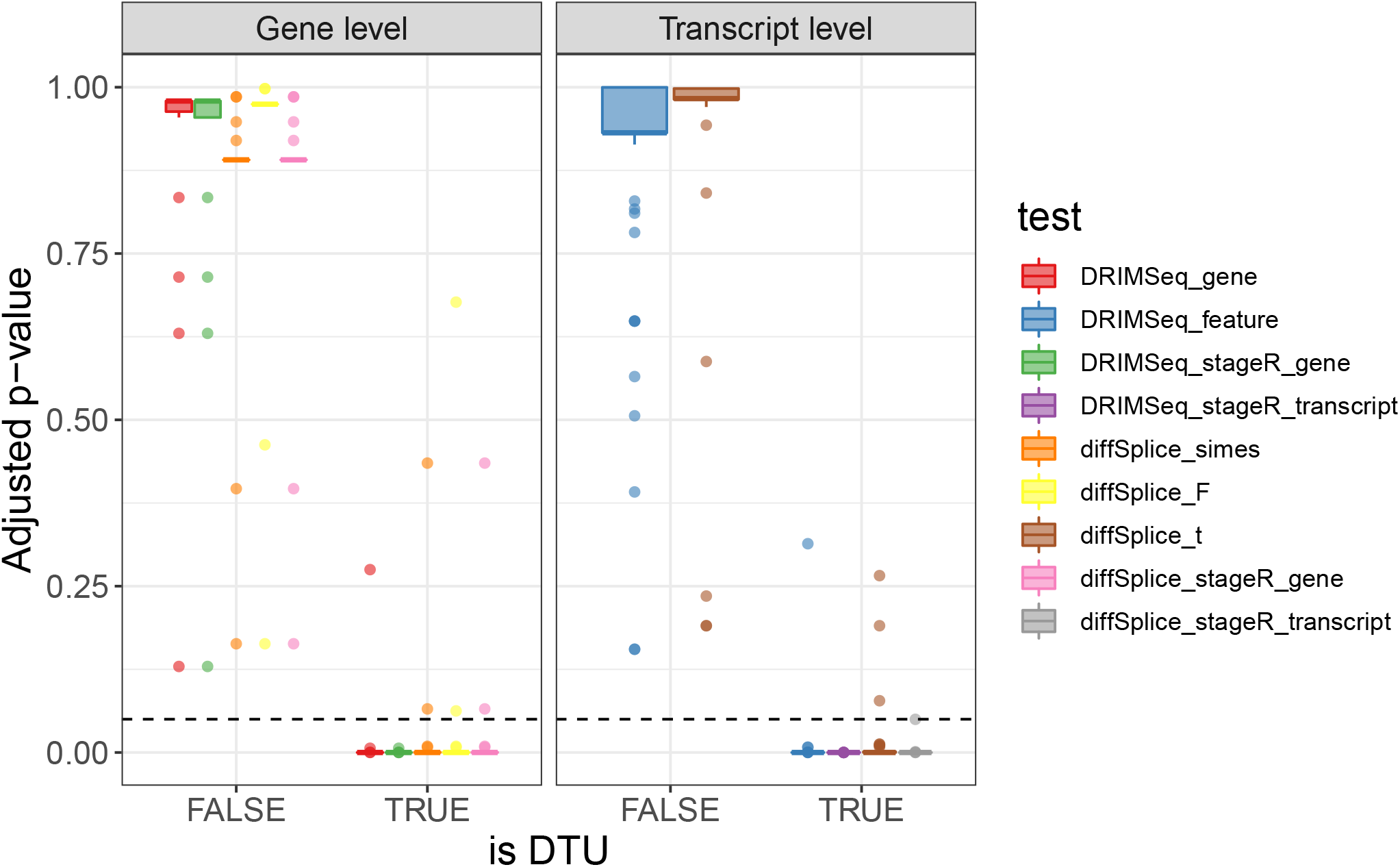
A box plot showing the distribution of adjusted *p*-values from different tests of DTU for the sequins data, faceted by whether the test is performed at the gene-level or transcript-level and stratified by whether the gene or transcript has true DTU. The dashed horizontal line shows the adjusted *p*-value cutoff of 0.05.

**Fig. S6.**
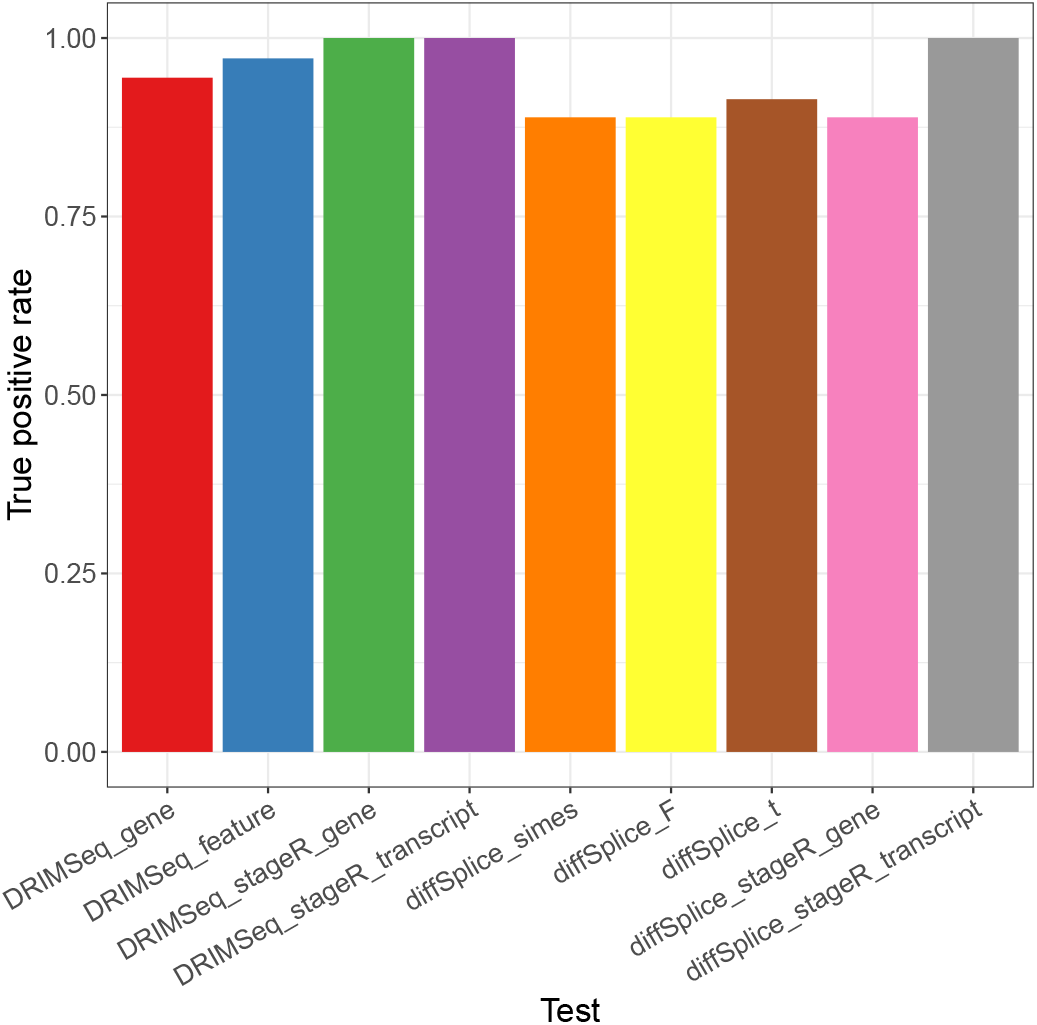
A bar plot showing the true positive rate (TPR) from different tests of DTU for the sequins data.

**Fig. S7.**
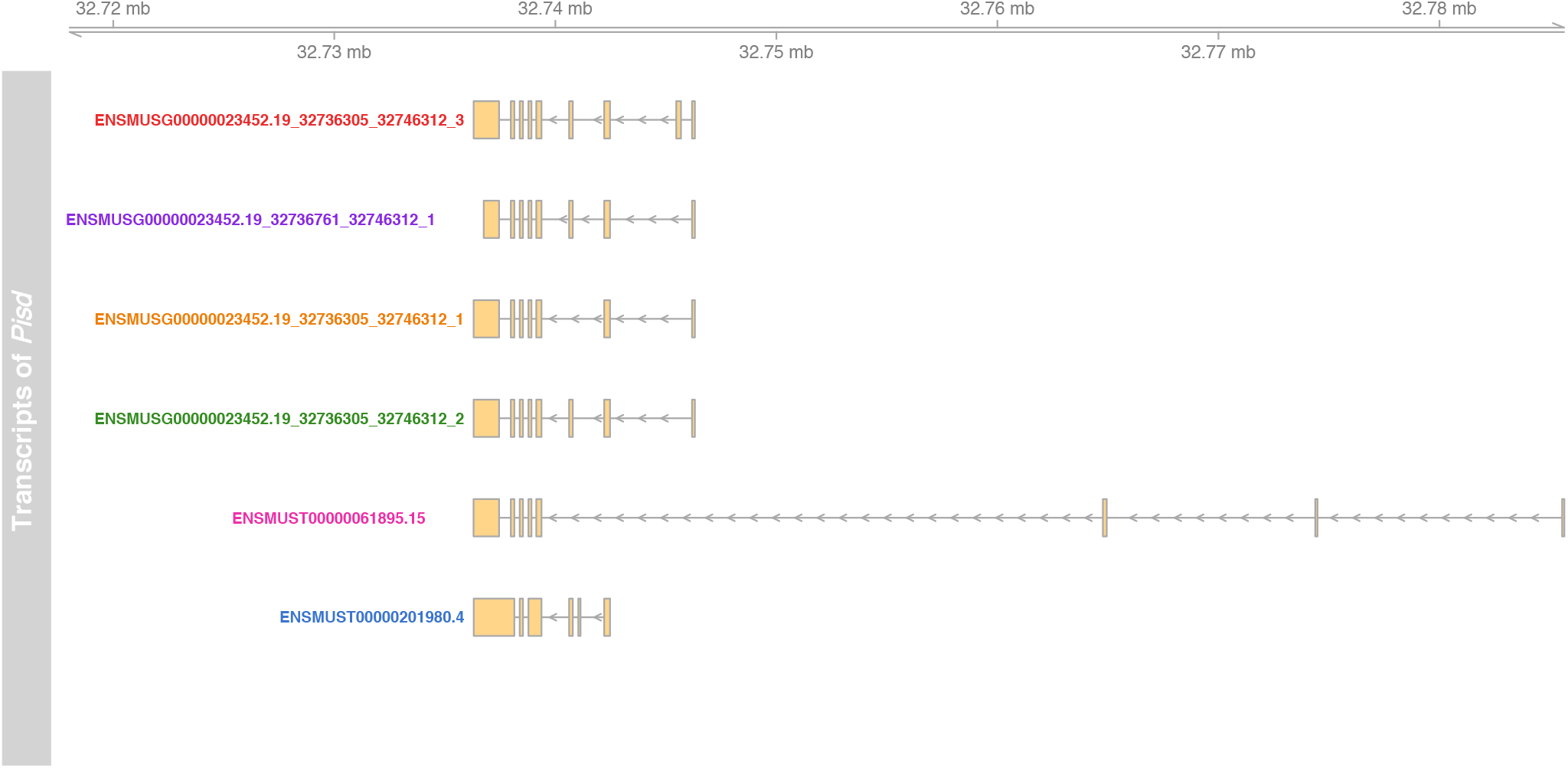
Different isoforms of gene *Pisd* in the NSC dataset identified by *FLAMES*.

**Fig. S8.**
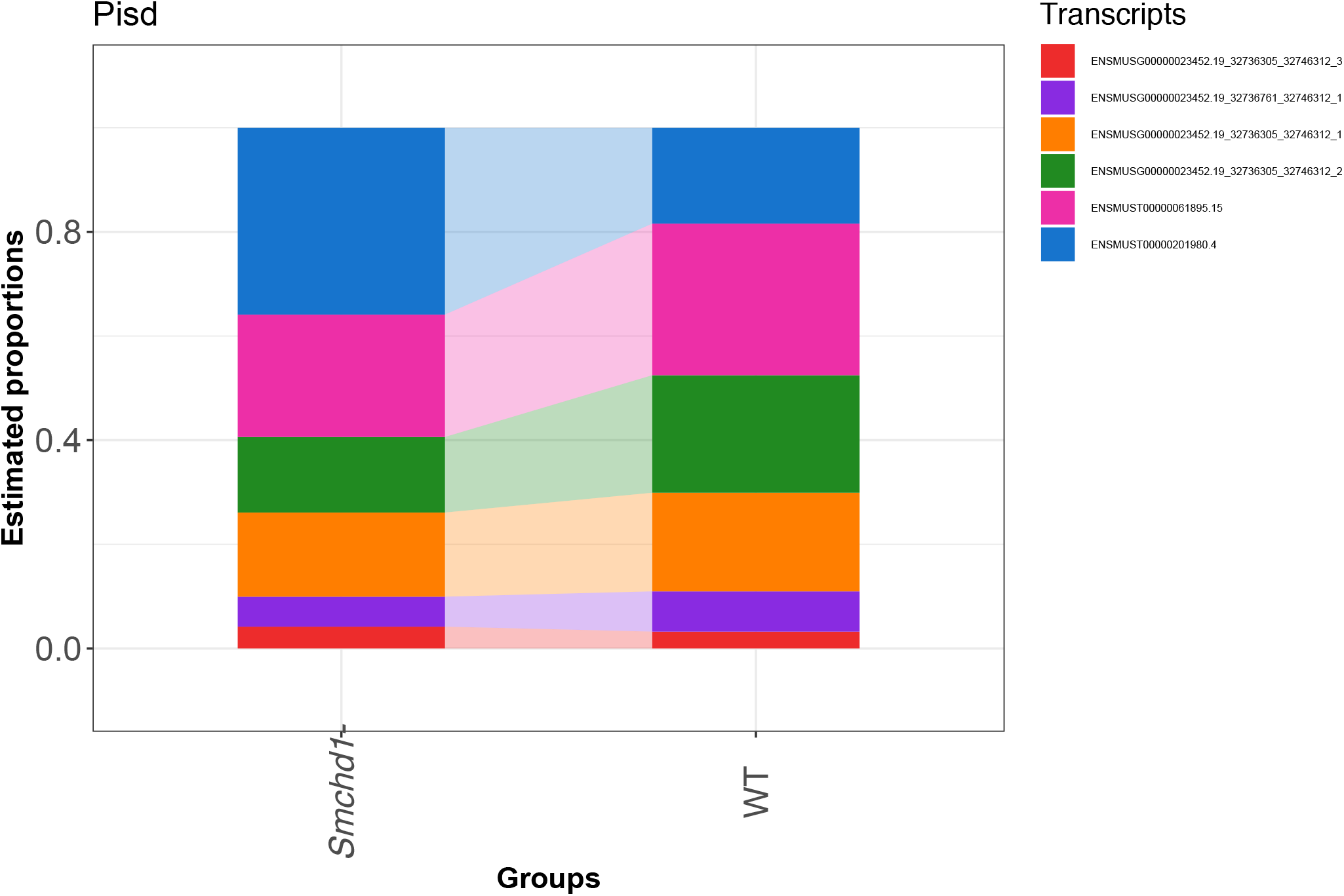
A ribbon plot showing the estimated isoform proportions for the gene *Pisd* in the NSC dataset.

## Notes

### Competing Interest Statement

MBC and RDP have received support from Oxford Nanopore Technologies (ONT) to present their findings at scientific conferences. However, ONT played no role in study design, execution, analysis or publication.

